# Chromosome fissions and fusions act as barriers to gene flow between *Brenthis* fritillary butterflies

**DOI:** 10.1101/2022.10.30.514431

**Authors:** Alexander Mackintosh, Roger Vila, Dominik R. Laetsch, Alex Hayward, Simon H. Martin, Konrad Lohse

## Abstract

Chromosome rearrangements are thought to promote reproductive isolation between incipient species. However, it is unclear how often, and under what conditions, fission and fusion rearrangements act as barriers to gene flow. Here we investigate speciation between two largely sympatric fritillary butterflies, *Brenthis daphne* and B. ino. We use a composite likelihood approach to infer the demographic history of these species from whole genome sequence data. We then compare chromosome-level genome assemblies of individuals from each species and identify a total of nine chromosome fissions and fusions. Finally, we fit a demographic model where effective population sizes and effective migration rate vary across the genome, allowing us to quantify the effects of chromosome rearrangements on reproductive isolation. We show that chromosomes involved in rearrangements experienced less effective migration since the onset of species divergence and that genomic regions near rearrangement points have a further reduction in effective migration rate. Our results suggest that the evolution of multiple rearrangements in the *B. daphne* and *B. ino* populations, including alternative fusions of the same chromosomes, have resulted in a reduction in gene flow. While fission and fusion of chromosomes are unlikely to be the only processes that have led to speciation between these butterflies, this study shows that these rearrangements can directly promote reproductive isolation and may be involved in speciation when karyotypes evolve quickly.

## Introduction

The process of speciation, where groups of individuals become reproductively isolated from one another, is driven by evolutionary forces that prevent gene flow. Many closely related species show differences in karyotype and there has been much discussion about the role of chromosome rearrangements (e.g. inversions, translocations, fissions, and fusions) in preventing gene flow and promoting speciation. Early work on *Drosophila* demonstrated that inversions suppress recombination (Sturtevant 1921; Dobzhansky and Epling 1948). More recently, both theoretical models (Navarro and Barton 2003; Kirkpatrick and Barton 2006) and examples in a variety of organisms (Wellenreuther and Bernatchez 2018) have shown that inversions can facilitate local adaptation, promote the evolution of genetic incompatibilities and act as barriers between recently diverged species. It is less clear, however, whether fission and fusion rearrangements have a similarly important role in speciation (Rieseberg 2001). These rearrangements do not typically confer the same drastic change in recombination as inversions do, yet there is evidence for increased speciation rates in groups where fissions and fusions happen more often (Bush *et al*. 1977; Leaché *et al*. 2016; de Vos *et al*. 2020). Fissions and fusions could act as barriers to gene flow if hybrid individuals that are heterozygous for a rearrangement suffer from underdominance (heterozygote disadvantage). This will happen when karyotypic heterozygosity generates multivalents at meiosis, which are prone to unbalanced segregation. While there is indeed evidence for fissions and fusions causing underdomi-nance through aneuploidy (Dutrillaux and Rumpler 1977; Castiglia and Capanna 2000; Lukhtanov *et al*. 2018), models of chromosomal speciation that assume underdominance are paradoxical; for hybrids to suffer from underdominance the rearrangement must be at high frequency in one population, but how does a rearrangement rise to high frequency if it causes underdominance? Proposed solutions to this paradox include fixation by meiotic drive (White 1968), strong drift in a founder population (Templeton 1981; but see Barton and Charlesworth 1984), and complex rearrangements that evolve in a stepwise manner, where each step has a small fitness effect (White 1978b; Baker and Bickham 1986). This limits the conditions under which underdominant chromosomal speciation can happen, and it is therefore perhaps unsurprising that there are few convincing empirical examples (see Basset *et al*. 2006 and Yannic *et al*. 2009).

Not all models of chromosomal speciation require underdominance. For example, fusions could affect gene flow by bringing pre-existing barrier loci onto the same chromosome. Guerrero and Kirkpatrick (2014) showed that for two polymorphic loci maintained by selection-migration balance, a fusion will rise in frequency if it brings two locally adapted alleles into strong linkage disequilibrium (LD). This process has the potential to strengthen the combined effect of barrier loci by reducing recombination between them, thus promoting reproductive isolation. Although Guerrero and Kirkpatrick (2014) do not include underdominance in their model, the process they describe is not mutually exclusive with underdominant chromosomal speciation, and may offer an additional way for fusions to evolve in spite of underdominance.

Fission and fusion rearrangements can also influence the accumulation of reproductive isolation when a barrier to gene flow is highly polygenic. Given such a barrier, the probability that a neutral allele migrates is partly determined by whether it can recombine away from the foreign deleterious alleles that it was introgressed with (Aeschbacher *et al*. 2017). Fissions and fusions can alter the per-base recombination rates of chromosomes by changing their length and they can therefore influence effective migration. Recently, Martin *et al*. (2019) showed that recombination rate was the main determinant of the amount of introgression between species of *Heliconius* butterflies, with long fused chromosomes having less introgression than short non-fused ones. These fusions cannot be barriers themselves because they are shared among the species. Instead, because of their length, the fused chromosomes have a low per-base crossover rate (Davey *et al*. 2017), which reduces effective migration when barrier loci are common. While the fusions in these *Heliconius* butterflies are shared, similar logic applies to a fusion that generates a long chromosome in just one population.

If a pair of species’ karyotypes differ by fissions and fusions, it is entirely possible that these rearrangements have not played any significant role in their speciation. Rearrangements can arise once reproductive isolation is already complete, meaning that other barriers must have driven speciation. Alternatively, if rearrangements are present during the early stages of speciation, they may not have any effect on gene flow. This would be the case if underdominance was weak enough for a rearrangement to be effectively neutral. Moreover, even if rearrangements do have underdominant or recombination modifying effects, there may be barriers of very large effect which have played a much greater role in speciation. It is therefore important to quantify the effect of fission and fusion rearrangements on gene flow, rather than assuming that these conspicuous changes in the genome must play an important role in the speciation process.

Most Lepidoptera (moths and butterflies) have similar karyotypes, consisting of around 30 pairs of autosomes and ZW sex chromosomes. However, there are notable exceptions. For example, *Pieris* butterflies have a reduced karyotype where chromosomes have undergone substantial reorganisation via inter-chromosomal rearrangements (Hill *et al*. 2019). There are also groups with highly variable chromosome counts, such as the butterfly genera *Erebia, Lysandra, Polyommatus*, and *Leptidea*. In each of these genera it has been suggested that rearrangements may have facilitated speciation (Talavera *et al*. 2013; Lukhtanov *et al*. 2005, 2011), although the extent to which rearrangements have affected gene flow remains unclear.

Another karyotypically variable group of butterflies is the genus *Brenthis* (Nymphalidae) which consists of four species. While 34 chromosome pairs have been observed in *Brenthis hecate* spermatocytes (de Lesse 1961; Saitoh and Lukhtanov 1988), *B. daphne* and *B. ino* are reported to have only 12 - 14 pairs of chromosomes (Maeki and Makino 1953; Saitoh *et al*. 1989; Federley 1938; Saitoh 1987, 1991). We recently assembled a *B. ino* reference genome (Mackintosh *et al*. 2022) with 14 pairs of chromosomes. We found that a male individual was heterozygous for a neo-Z chromosome fusion and that the genome was highly rearranged compared to the ancestral nymphalid karyotype. These results are consistent with rapid, and likely still ongoing, chromosome evolution in the genus *Brenthis*.

The sister species *B. daphne* and *B. ino* are largely sympatric (Figure 1), have differences in larval host plant preference, and are estimated to have split approximately 3 million years ago (Ebdon *et al*. 2021). Interspecific mating experiments have shown that female *B. daphne* and male *B. ino* can produce fertile offspring, suggesting that reproductive isolation between these species is incomplete (Kitahara 2008, 2012). Additionally, putative F1 hybrids have been observed in Japan (Kitahara 2012). Similar chromosome numbers have been observed for males of either species, 12 - 14 for *B. daphne* and 13 - 14 for *B. ino*, suggesting intraspecific variation in karyotype, but no large differences between species. However, chromosome numbers will be unchanged by reciprocal translocations or an equal number of chromosome fission and fusion events. Such ‘cryptic’ rearrangements are best identified by comparing genome assemblies. If *B. daphne* and *B. ino* possess cryptic inter-chromosomal rearrangements, then their recent divergence and potential for ongoing gene flow makes them a useful model for investigating the effects of rearrangements on reproductive isolation.

**Figure 1:**
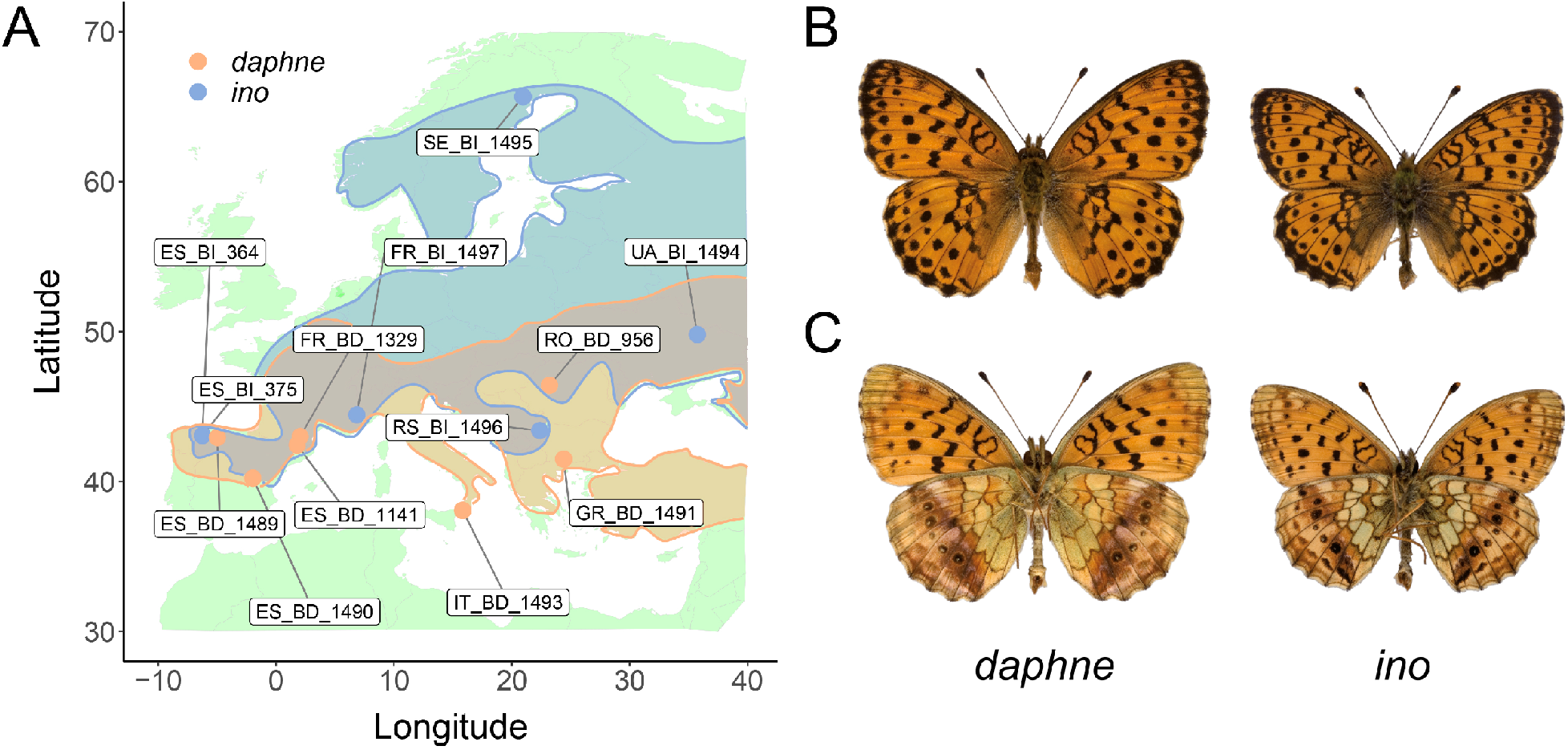
(A) Sampling locations of *Brenthis daphne* (orange) and *B. ino* (blue) individuals across Europe. Approximate distributions are also shown using the same colour scheme. (B) Uppersides of male *B. daphne* and male *B. ino*. (C) Undersides of male *B. daphne* and male *B. ino*.

Here we show that the genomes of *B. daphne* and *B. ino* differ in synteny due to multiple fission and fusion rearrangements. More specifically, almost half of the chromosomes are involved in rearrangements whereas the rest are syntenic. We estimate the demographic history of these species as well as genome-wide variation in effective migration rate (*m_e_*). By intersecting estimates of *m_e_* with chromosome rearrangements, we test whether fissions and fusions have acted as barriers to gene flow. We consider the following scenarios:

- **No effect:** Fission and fusion rearrangements are selectively neutral and have had no effect on the effective rate of gene flow, either directly or indirectly.
- **Underdominance:** Fissions and fusions produce direct, localised barriers to gene flow because early generation hybrids and backcrosses with heterokaryotypes have reduced fitness.
- **Fused barriers:** Fusions are not barriers to gene flow themselves, but have brought individual barrier alleles of large effect into linkage, thus strengthening the barrier effect of these loci.
- **Polygenic barriers:** In the presence of polygenic barriers, fissions and fusions affect gene flow by modifying chromosome lengths and therefore recombination rates.

## Results

### Diversity and divergence

Using our previously published *B. ino* genome assembly (Mackintosh *et al*. 2022) as a reference, we analysed whole genome sequence data for seven *B. daphne* and six *B. ino* individuals (Figure 1; Table S1). We restricted our analyses to intergenic regions of the genome, as these typically evolve under less selective constraint than genic regions. Consistent with a previous analysis of transcriptomic data (Ebdon *et al*. 2021), we find that per-site heterozygosity is greater in *B. ino* (0.0111) than in *B. daphne* (0.0043) and that interspecific divergence is considerable (d_xy_ = 0.0228, F_ST_ = 0.4976). We also find evidence of shallow population structure within each species (Figures 2A and 2B). For example, pairwise F_ST_ is ~ 0.1 for *B. daphne* individuals sampled in different glacial refugia (Iberia, Italy, or the Balkans) and there are similar levels of differentiation between *B. ino* individuals sampled from Iberia and elsewhere in Europe. While this shows that European *B. daphne* and *B. ino* are not panmictic populations, this should only have a small effect on our analyses of long-term divergence and gene flow between the two species (see below).

**Figure 2:**
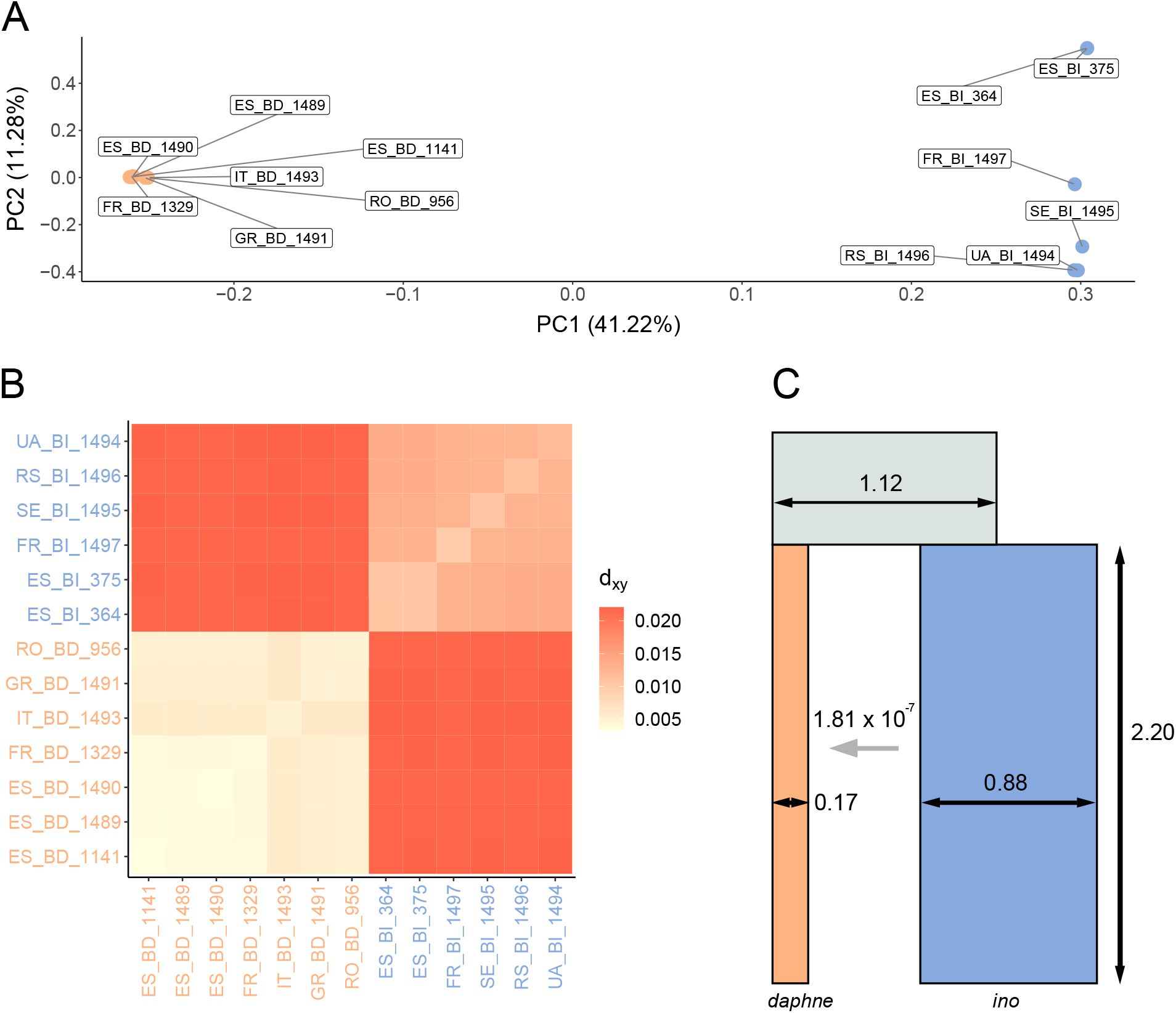
Diversity and divergence between *B. daphne* and *B. ino*. (A) A PCA of individuals sampled across Europe, with PC1 corresponding to interspecific variation. Orange points are *B. daphne* individuals and blue points are *B. ino* individuals. The same colour scheme is used in subplots (B) and (C). (B) A heatmap showing *d_xy_* between pairs of individuals. (C) The best fitting demographic model, with parameter values inferred from the genome-wide bSFS. The *N_e_* (indicated by horizontal black arrows) and split time (vertical black arrow) parameters are in units of 10^6^. The horizontal grey arrow indicates the direction of gene flow, from *B. ino* to *B. daphne*, forwards in time.

### Demographic history

We use gIMble (Laetsch *et al*. 2022), a recent implementation of a blockwise likelihood calculation (Lohse *et al*. 2016), to infer the demographic history of speciation between *B. daphne* and *B. ino*. gIMble calculates the blockwise site frequency spectrum (bSFS) of all possible interspecific pairwise comparisons, i.e. sampling a single diploid genome from each species and tallying mutations in short blocks of sequence (see Methods). We fit three demographic models to the bSFS with gIMble: strict divergence (*DIV*) and two scenarios of isolation with migration (*IM*_→*Bda*_ and *IM*_→*Bin*_). The DIV model has three *N_e_* parameters (*B. daphne, B. ino*, ancestral) and a split time parameter. The IM models have an additional parameter, i.e. they assume a constant rate of effective migration (*m_e_*) either from *B. ino* into *B. daphne* forwards in time (*IM*_→*Bda*_) (Figure 2C) or in the opposite direction (*IM*_→*Bin*_). By optimising the parameters within each model, we found that the *IM*_→*Bda*_ model fits best (Table 1; Figure 2C). The *DIV* and *IM*_→*Bin*_ models converged to the same parameter values and composite likelihood (Table 1), i.e. the maximum composite likelihood (MCL) estimate of *m_e_* under the *IM*_→*Bin*_ model is 0. By contrast, the MCL estimate of *m_e_* from *B. ino* to *B. daphne* within the best fitting (*IM*_→*Bda*_) model is 1.811 × 10^-7^, which is equivalent to 0.124 effective migrants per generation. As a result of this migration, the *IM*_→*Bda*_ model also has a older split time (≈ 2.2 MY) than the *DIV/IM*_→*Bin*_ model (≈ 1.2 MY) (Table 1).

**Table 1:** Maximum composite likelihood parameters for three different demographic models. The *IM*_→*Bda*_ model has the greatest lnCL.

| Model | $N_e$ *daphne* | $N_e$ *ino* | $N_e$ *ancestral* | $m_e$ | Split time | $\ln CL$ |
| --- | --- | --- | --- | --- | --- | --- |
| *DIV* | 252,395 | 682,995 | 1,432,643 | - | 1,182,959 | -234,678,837 |
| $IM_{\rightarrow Bda}$ | 170,577 | 880,037 | 1,116,093 | $1.811 \times 10^{-7}$ | 2,201,678 | -233,968,576 |
| $IM_{\rightarrow Bin}$ | 252,395 | 682,995 | 1,432,643 | 0.000 | 1,182,958 | -234,678,837 |

An *IM* model has to fit the data equally well or better than a *DIV* model because it includes an additional parameter, *m_e_*. To test whether the *IM*_→*Bda*_ model fits significantly better than *DIV* (see Laetsch *et al*. 2022), we simulated parametric bootstrap replicates for the MCL estimates under the *DIV* history and optimised both the *DIV* and *IM*_→*Bda*_ models. The improvement in fit (Δ lnCL) between *DIV* and *IM*_→*Bda*_ models for parametric bootstrap replicates was far below what we observe in the data (Figure S2). An IM demographic history, with migration from *B. ino* to *B. daphne*, is therefore well supported.

### Synteny

To compare synteny between *B. daphne* and *B. ino*, we generated a chromosome-level assembly for a female *B. daphne* individual, collected in Catalunya, Spain. The assembly is 419.1 Mb in length, with a scaffold N50 of 30.6 Mb and a contig N50 of 13.4 Mb. The *B. daphne* assembly is scaffolded into 13 chromosome-level sequences (hereafter simply referred to as chromosomes) corresponding to 12 autosomes and the Z sex chromosome (Figure 3). We failed to scaffold the W chromosome which is likely contained within the remaining 35 contigs that total 5.3 Mb.

**Figure 3:**
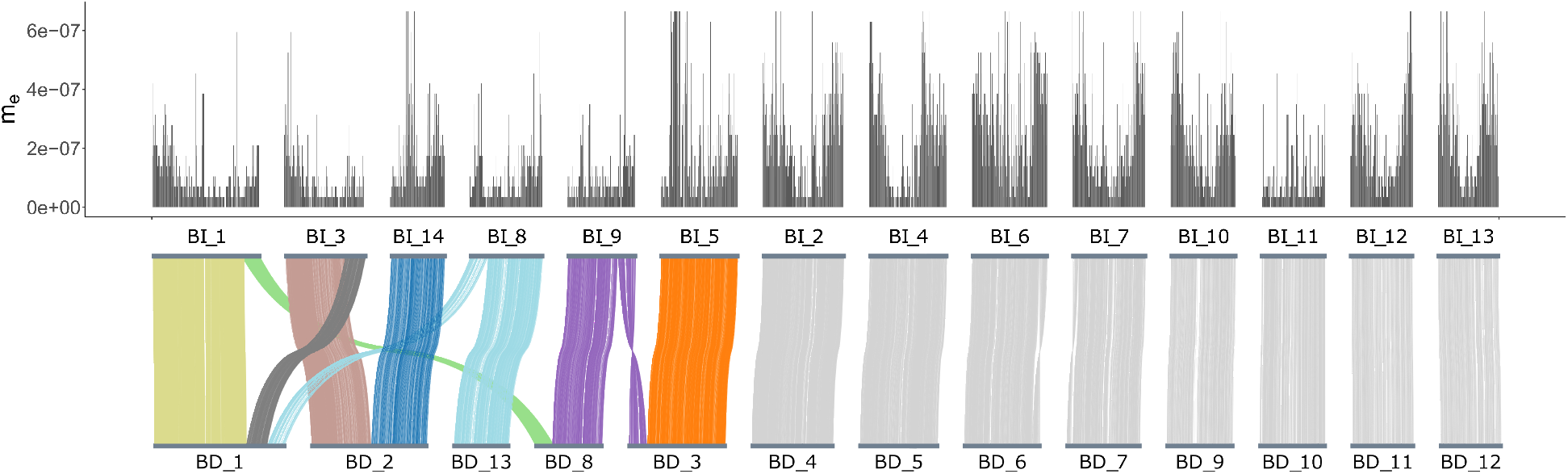
A whole genome alignment between *B. daphne* and *B. ino*, with effective migration (*m_e_*) estimates for windows along the *B. ino* genome plotted above. Alignments between non-rearranged chromosomes are coloured in grey. Alignments between rearranged chromosomes are coloured by the inferred chromosomes of the common ancestor of *B. daphne* and *B. ino*. The Z chromosome is labelled as BD_10 in the *B. daphne* genome and Bl_11 in the *B. ino* genome.

A pairwise alignment between the *B. daphne* and *B. ino* assemblies shows that only eight chromosomes have one-to-one homology, with the others showing more complex relationships (Figure 3). For example, *B. daphne* chromosome 1 is homologous to parts of *B. ino* chromosomes 1, 3, and 8 (Figure 3). Altogether, we find that five *B. daphne* chromosomes and six *B. ino* chromosomes are involved in a total of nine inter-chromosomal rearrangements. Hereafter we refer to these chromosomes as rearranged. Additionally, we define rearrangement points as chromosome ends involved in fissions / fusions or sites where alignments on either side connect different *B. daphne* and *B. ino* chromosomes.

From a single pairwise comparison it is not possible to tell whether a genome possesses a rearrangement in the ancestral or derived state. Therefore, to polarise these rearrangements, we analysed the assemblies alongside a publicly available genome assembly of *Fabriciana adippe* (see Methods). We infer a maximally parsimonious history of rearrangements where the common ancestor of *B. daphne* and *B. ino* had 16 chromosomes, with two fissions and five fusions in the *B. daphne* lineage and two fusions in the *B. ino* lineage. This inferred rearrangement history involves two small ancestral chromosomes (approximately 6.6 and 8.4 Mb), which fused independently to different chromosomes in either species (Figure 3).

### Variation in *m_e_* across the genome

To investigate the effect of rearrangements on reproductive isolation, we followed the approach of Laetsch *et al*. (2022) by inferring effective population sizes (*N_e_*) and the effective migration rate (*m_e_*) in windows along the genome. We assume that the species split time is fixed to the MCL estimate under the *IM*_→*Bda*_ model (Table 1). We used simulations to confirm that, given plausible (but conservative) assumptions about recombination, demographic parameters could be inferred for windows containing 30,000 consecutive sequence blocks (Supplementary note 1). To infer parameters for the real data, we set up a grid of 67,500 possible parameter value combinations: 15 *B. daphne N_e_* values (20,000 - 720,000), 15 *B. ino N_e_* values (50,000 - 2,850,000), 15 ancestral *N_e_* values (50,000 - 2,010,000), and 20 *m_e_* values (0 - 6.65 × 10^-7^). We identified the best fitting parameter combination for each window (30,000 consecutive blocks, median length = 122 kb). Estimates of local *m_e_* have a long tailed distribution with a peak at 3.5 × 10^-8^ (Figure 4). Consistent with the genome-wide estimate, the mean *m_e_* across windows is 1.845 × 10^-7^. We find that *m_e_* is lower on rearranged chromosomes compared to non-rearranged chromosomes (mean *m_e_* = 1.281 × 10^-7^ vs 2.292 × 10^-7^ respectively; Figure 3; Figure 4; permutation test p < 0.005). This suggests that inter-chromosomal rearrangements are associated with reduced gene flow.

**Figure 4:**
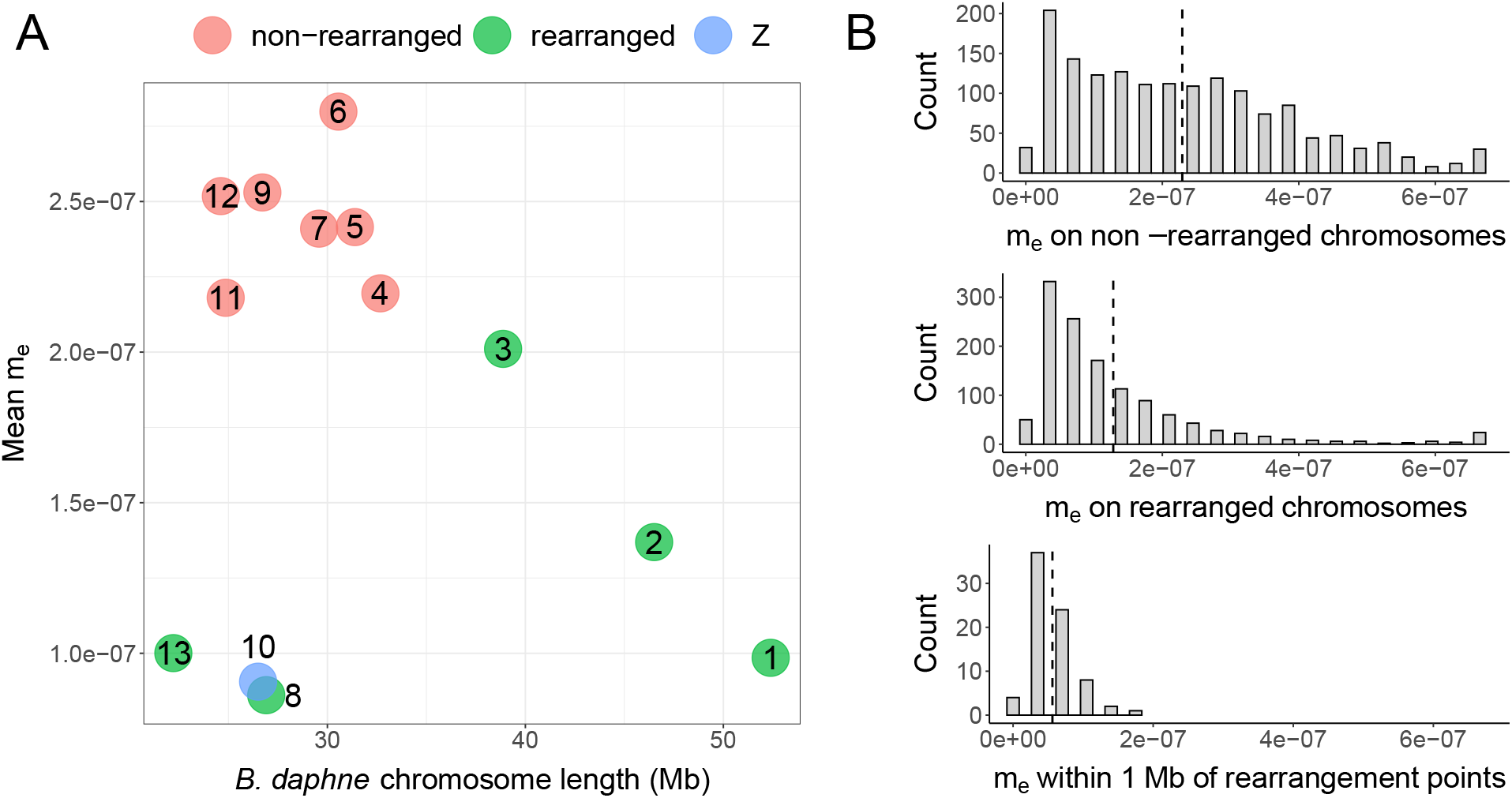
Differences in effective migration (*m_e_*) between rearranged and non-rearranged chromosomes. (A) Mean *m_e_* for each *B. daphne* chromosome plotted against its length. Points are coloured green if the chromosome is rearranged and red if not. The Z chromosome, which is not rearranged, is coloured blue. (B) The distribution of *m_e_* estimates across non-rearranged chromosomes (top), rearranged chromosomes (middle), and within regions near rearrangement points (bottom). For each plot, the mean is plotted as a dashed vertical line.

An alternative approach to estimating *m_e_* for each window is to identify ‘barrier windows’ where there is statistical support for a reduction in gene flow (compared to the background *m_e_*). Following Laetsch *et al*. (2022), we defined barrier windows as those where *m_e_* = 0 has a greater lnCL than *m_e_* = 1.75 × 10^-7^ (the grid value nearest to the genome-wide *m_e_* estimate). Under this criterion, 23.08% of windows are barriers and these are distributed across all 14 *B. ino* chromosomes. However, the number of barrier windows is not equal among *B. ino* chromosomes, e.g. 48.11% and 4.22% of windows are barriers on chromosome 3 and chromosome 10, respectively. Windows on rearranged chromosomes are twice as often classified as barriers than windows on non-rearranged chromosomes (32.91% vs 15.27%; permutation test p < 0.01). The window with the greatest barrier support (Δ lnCL) is located on *B. ino* chromosome 8, with the start of this window being less than 200 kb from a rearrangement point. This alternative, but not independent, estimation of *m_e_* variation provides further evidence for an association between fission and fusion rearrangements and a reduction in gene flow.

In the best fitting demographic model (Figure 2C) *B. daphne* receives gene flow from *B. ino*. As a result, low recombination regions in the *B. daphne* genome are expected to have reduced *m_e_* under the *polygenic barriers* scenario (see Introduction). With this in mind, it is therefore possible that our observation of reduced *m_e_* on rearranged chromosomes is not because of rearrangements acting as barriers directly, but is instead a result of rearrangements producing large *B. daphne* chromosomes with low recombination rates (e.g. *B. daphne* chromosomes 1, 2, and 3; see Figure 3). To test this possibility, we assigned each genomic window to a *B. daphne* chromosome using a whole genome alignment (Figure 3) and calculated the mean *m_e_* of each *B. daphne* chromosome. There is no significant linear relationship between *B. daphne* chromosome length and mean *m_e_* (Spearman’s _*pdf* =11_ = −0.0769, p = 0.8065; Figure 4). While the largest chromosomes, which happen to be rearranged, do indeed have relatively low *m_e_*, short rearranged chromosomes also have low *m_e_*. Additionally, the Z chromosome (*B. daphne* chromosome 10, *B. ino* chromosome 11), which is not rearranged and is short, has low mean *m_e_*. Chromosome size alone is therefore unlikely to explain the association between chromosome rearrangements and reduced *m_e_*.

If fission and fusion rearrangements act as direct barriers to gene flow, such as in the *fused barriers* and *underdominance* scenarios, then we would expect loci that are closely linked to rearrangement points to have the greatest reduction in *m_e_*. This is because loci that are on the same chromosome but are less closely linked will be more likely to recombine away following introgression. Selection against foreign rearrangements will therefore only have a weak effect on loosely linked loci. We indeed find that genomic windows which are located within 1 Mb of a rearrangement point have a lower m_e_ (mean = 5.618 × 10^-8^) than those located elsewhere on rearranged chromosomes (mean = 1.328 × 10^-7^) (Figure 4; permutation test p < 0.0005. All 76 of these windows have estimated *m_e_* values (between 0 and 1.75 × 10^-7^; Figure 4) that are below the genome-wide estimate (1.811 × 10^-7^). Additionally, 59.21% of them are classified as barrier windows. The signal of reduced *m*e at closely linked sites provides support for rearrangements having acted as barriers to gene flow.

## Discussion

We have shown that the fritillary butterflies *Brenthis daphne* and *B. ino* possess different karyotypes due to multiple fission and fusion rearrangements, and that these rearrangements are associated with reduced *m_e_*. We can therefore reject the *no effect* scenario where rearrangements are only coincidental with speciation.

We considered the possibility that the association between rearrangements and low *m_e_* could be driven by the modification of chromosome lengths, and therefore recombination rate, in the presence of polygenic barriers. Indeed fusions in the *B. daphne* population have generated large (up to 52 Mb) chromosomes with presumably low recombination rates and low *m_e_*. While recombination rate certainly plays a role in determining variation in *m_e_* across the genome (see below), the fact that small chromosomes that are involved in fissions and fusions have reduced *m_e_* (Figure 4) is not well explained by the *polygenic barriers* scenario where rearrangements only modify the size of chromosomes. Our results are therefore best explained by fissions and fusions leading to localised selection against introgression around the rearrangement points.

The association between rearrangements and *m_e_* that we find is consistent with two scenarios, *underdominance* and *fused barriers*. Under the *underdominance* scenario we would expect rearranged chromosomes to have lower *m_e_* and we would also expect *m_e_* to be further reduced near rearrangement points. We find both of these patterns in our data (Figure 4). The expectations under the *fused barriers* scenario are more variable. If the number of initial barrier loci is small, and fusions that put two or more barrier loci into strong LD rise in frequency due to natural selection (Guerrero and Kirkpatrick 2014), then we would indeed expect lower *m_e_* on rearranged chromosomes as well as particularly low *m_e_* around fusion points. However, if there were enough initial barrier loci so that some were in tight linkage by chance alone, then the *m_e_* of barrier loci brought together by a fusion would be unremarkable. One way to discern between the *fused barriers* and *underdominance* scenarios would be to compare *m_e_* around fission points, as it is only expected to be reduced in the *underdominance* scenario. However, the two fission events in the *B. daphne* lineage are both followed by fusions, making this test inappropriate. So while the *fused barriers* scenario requires a particular number and distribution of initial barrier loci, it is still consistent with our results. Note, again, that the *fused barriers* and *underdominance* scenarios are not mutually exclusive, and both processes could have contributed to fissions and fusions acting as barriers to gene flow between *B. daphne* and *B. ino*.

Earlier we noted that chromosomal speciation models involving underdominance are often paradoxical (see Introduction). So, how could rearrangements rise to high frequency in the *B. daphne* and *B. ino* populations if heterokaryotypes are selected against? The *fused barriers* scenario is one way in which underdominance could be overcome within a population, although it can only explain the evolution of fission rearrangements if they are translocations instead. Another solution is that the fitness consequences of heterozygosity for a single fission / fusion are effectively neutral. This is more likely to be the case when chromosomes are holocentric (Lucek *et al*. 2022), as they are in butterflies (although see Dutrillaux *et al*. 2022). A single rearrangement could therefore fix in a population and, over time, karyotypes could evolve in a stepwise process. By contrast, heterozy-gosity for multiple fissions / fusions could have a larger fitness cost due to the difficulty of properly segregating multiple, potentially complex, multivalents (Dutrillaux and Rumpler 1977; Castiglia and Capanna 2000). If *B. daphne* and *B. ino* evolved multiple rearrangements through a stepwise process during a period of allopatry, then rearrangements could act as barriers once the populations came back into contact. This scenario, which has similarities with the stepwise accumulation of Bateson–Dobzhansky–Muller incompatibilities (Dobzhansky 1934), has been previously described by White (1978a), and is known as the chain model (Rieseberg 2001). While the rearrangements between *B. daphne* and *B. ino* are numerous and complex (Figure 3), consistent with the chain model, we have not tested whether these populations underwent a period of allopatry followed by secondary contact. There may be enough information in the two-diploid bSFS to fit such a model, but no exact likelihood implementation exists yet (although see Bisschop 2022 and Beeravolu *et al*. 2018). Importantly, if the chain model does apply here, it has only generated partial barriers to gene flow and has not resulted in complete reproductive isolation. If hybrids with heterokaryotypes were sterile, then gene flow would cease across the entire genome. We instead find that gene flow is reduced on rearranged chromosomes, which means that heterokaryotype hybrids must have been able to backcross.

We have focused on whether chromosome rearrangements, the most conspicuous genomic difference between these species, have acted as barriers to gene flow. Yet variation in *m_e_* across the genome cannot be explained by rearrangements alone. Firstly, the centres of non-rearranged chromosomes clearly have lower *m_e_* estimates than regions near chromosome ends (Figure 3). This can be explained by variation in recombination rate, with crossovers concentrated towards telomeres (Haenel *et al*. 2018), as neutral alleles are more likely to introgress if they can quickly recombine away from the barrier loci they are linked to. The fact that chromosome centres consistently have lower *m_e_* suggests that there are other barriers to gene flow distributed across the genome, not only rearrangement points. Secondly, the Z chromosome has a considerably lower mean *m_e_* than all other non-rearranged chromosomes (Figure 4), which cannot be because of rearrangements or low recombination (the Z recombines more frequently than autosomes in Lepidoptera). Instead, low *m_e_* on the Z may be a result of recessive barrier loci being exposed to selection in females (Turelli and Orr 1995). Additionally, if the Z evolves faster than autosomal chromosomes (Mongue *et al*. 2021) then barrier loci, both recessive and dominant, may accumulate faster. Reduced gene flow on the *Brenthis* Z chromosome mirrors findings in other butterfly systems (Xiong *et al*. 2022; Rosser *et al*. 2022), as well as in birds (Irwin 2018; Ottenburghs 2022), suggesting that Z chromosomes often accumulate reproductive isolation at a faster rate than autosomes. Given that there are likely many barriers to gene flow between *B. daphne* and *B. ino*, especially on the Z, it may be inaccurate to describe the history of these species as ‘chromosomal speciation’. Instead, fission and fusion rearrangements are likely one of several processes that have promoted reproductive isolation.

The particular process we have investigated here, where fissions and fusions act as barriers to gene flow, likely modulates speciation more strongly in certain groups of organisms than in others. For example, the majority of butterfly species have very slow karyotypic evolution and thus speciation will have happened through the accumulation of other genetic barriers. Nevertheless, radiations of butterflies where karyotypes evolve quickly (e.g. the genera *Erebia, Lysandra*, and *Polyommatus*) may be partly explained by fissions and fusions acting as barriers to gene flow. This could also be true for other karyotypically variable radiations, such as Rock-wallabies (Potter *et al*. 2017), Morabine grasshoppers (White *et al*. 1964; Kawakami *et al*. 2011), and Carex sedges (Márquez-Corro *et al*. 2021). Evidence for fissions and fusions promoting speciation has often been macro-evolutionary, where analyses of large phylogenetic trees have shown an association between rearrangement and diversification rates. By contrast, focusing on a single pair of species, we have shown that fissions and fusions can act as barriers to gene flow and that their effect can be quantified from genomic data.

## Materials and methods

### Sampling

Butterflies were collected by hand netting. Individuals collected by KL were flash frozen in a liquid nitrogen dry shipper (Table S1). Those collected by RV were dried and, after some days, stored in ethanol at −20°C (Table S1).

### Sequencing

Previously published data - the *B. ino* genome assembly and whole genome sequencing (WGS) data from three individuals (NCBI accessions: GCA_921882275.1; ERX7241006; ERX7249694; ERX7250096) - were used in this study (Table S1). The sequencing process for generating these data is described in Mackintosh *et al*. (2022). Additional sequence data - Pacbio long reads, HiC data, and WGS data for ten individuals - were generated for this study (Table S1).

A high molecular weight (HMW) DNA extraction was performed for *B. daphne* individual ES_BD_1141 (Table S1), using a salting-out protocol (see Mackintosh *et al*. 2022 for details). A SMRTbell sequencing library was generated from the HMW extraction by the Exeter Sequencing Service. This was sequenced on three SMRT cells on a Sequel I instrument to generate 20.4 Gb of Pacbio continuous long read (CLR) data.

A second *B. daphne* individual (FR_BD_1329; Table S1) was used for chromatin conformation capture (HiC) sequencing. The HiC reaction was done using an Arima-HiC kit, following the manufacturer’s instructions for flash frozen animal tissue. The Illumina TruSeq library was sequenced on an Illumina NovaSeq 6000 at Edinburgh Genomics, generating 9.9 Gb of paired-end reads.

DNA extractions were performed for nine individuals using a Qiagen DNeasy Blood & Tissue kit, following the manufacturers instructions. TruSeq Nano gel free libraries were prepared from these extractions as well as the HMW extraction of individual ES_BD_1141. All ten libraries were sequenced on a NovaSeq 6000 at Edinburgh Genomics, generating between 10.1 and 40.0 Gb of paired-end reads for each sample.

### Genome assembly

A *B. daphne* genome sequence was assembled from the Pacbio long reads (ES_BD_1141), HiC data (FR_BD_1329), and WGS data (ES_BD_1141) using the same pipeline described in Mackintosh *et al*. (2022) (Hu 2021; Aury and Istace 2021; Laetsch and Blaxter 2017; Guan *et al*. 2020; Durand *et al*. 2016; Robinson *et al*. 2018), with one modification; YaHS (Zhou *et al*. 2022) was used to scaffold the contig assembly into chromosomes rather than 3d-dna (Dudchenko *et al*. 2017).

### Synteny analysis

To identify rearrangements, the *B. daphne* and *B. ino* assemblies were aligned with minimap2 v2.17 (Li 2018) using the option -x asm10. Alignments longer than 50 kb and with a mapping quality of 60 were visualised with minimap2synteny.py.

To polarise rearrangements and infer ancestral chromosomes, the *Brenthis* assemblies were analysed alongside a *Fabriciana adippe* genome assembly (NCBI accession: GCA_905404265.1). Single copy orthologues were identified in each genome using BUSCO v5.3.2 (Simão *et al*. 2015) with the lepidoptera_odb10 dataset. Complete and Fragmented BUSCO genes were analysed with syngraph (https://github.com/DRL/syngraph). In brief, syngraph identifies sets of markers, in this case BUSCO genes, that are found on the same chromosome in all three assemblies. Which sets of markers are found together on extant chromosomes is also recorded. Then, given a phylogenetic tree, parsimony is used to estimate the marker content of ancestral chromosomes and the inter-chromosomal rearrangements on each branch.

### Variant calling and filtering

Raw WGS reads were adapter and quality trimmed with fastp v0.2.1 (Chen *et al*. 2018) and aligned to the *B. ino* assembly (GCA_921882275.1) with bwa-mem v0.7.17 (Li 2013). Duplicates were marked using sambamba v0.6.6 (Tarasov *et al*. 2015). Variants were called with freebayes v1.3.2-dirty (Garrison and Marth 2012), using the following options: -limit-coverage 250 -use-best-n-alleles 8 -no-population-priors -use-mapping-quality -ploidy 2 -haplotype-length -1.

Variant calls were filtered using gIMble preprocess (Laetsch *et al*. 2022), with the following options: -snpgap 2 -min_qual 10 -min_depth 8 -max_depth 3, where -max_depth is in units of mean coverage.

Callable sites for each individual were identified with mosdepth v0.3.2 (Pedersen and Quinlan 2017), called through gIMble preprocess. To restrict downstream analyses to intergenic regions of the genome, the callable sites bed file was stripped of sites belonging to genic and/or repeat regions.

### Summaries of diversity and divergence

Variants in intergenic regions of autosomal chromosomes, where all individuals had a genotype, were used to generate a PCA with plink v1.90b6.18 (Purcell *et al*. 2007).

Pairwise statistics (*d_xy_* and *F_ST_*) were calculated from the same set of variants using VCF_stats.py. The denominator for *d_xy_* was the total number of autosomal intergenic sites that were callable across all individuals (123 Mb out of a possible 150 Mb).

### Demographic modelling with gIMble

To fit a genome-wide demographic model, autosomal variants were analysed with gIMble. Blocks of 64 bases, with a max span of 128 bases, were generated for all interspecific pairwise comparisons. A bSFS with a *k_max_* values of 2 was tallied from these blocks. The bSFS contains mutation counts for 81,104,834 interspecific blocks, each of length 64 bases, distributed over 139 Mb of intergenic sequence. Three models (SI, *IM*_→*Bda*_, *IM*_→*Bin*_) were fit to the genome-wide bSFS and the model with the highest lnCL (*IM*_→*Bda*_) was used for downstream analysis. Absolute parameter estimates were calculated by assuming the *de novo* mutation rate estimate for *Heliconius melpomene* (2.9 × 10^-9^ mutations per site per generation; Keightley *et al*. 2015) and a generation time of one year.

Parametric bootstrap simulations were performed with msprime v1.0.2 (Baumdicker *et al*. 2021), called through gIMble simulate. The simulations were parameterised with the maximum composite likelihood (MCL) *DIV* values, i.e. the best fitting history without gene flow, and a per-base recombination rate of 8.5 × 10^-9^ (equivalent to a single crossover per male meiosis for 14 chromosome pairs). A total of 100 replicates were performed. Each simulated bSFS was optimised under the *DIV* and *IM*_→*Bda*_ models.

To estimate variation in *m_e_* and N*e* across the genome, genomic windows containing 30,000 consecutive blocks were defined. Next, likelihood calculations were generated for a grid of 67,500 parameter combinations using gIMble makegrid. The lnCL of each windowed bSFS was then calculated for every grid-point. The MCL grid-point was recorded for each window. Additionally, MCLs were recorded for each window conditioning on each *m_e_* value, e.g. the MCL of a window considering all grid-points where *m_e_* = 0.

Variation in *m_e_* across the Z chromosome was estimated as above, with the following modification: only male individuals (two *B. daphne*, three *B. ino*, Table S1) were analysed (since females only have a single copy of the Z). Given the smaller number of interspecific comparisons (6 vs 42 for the autosomal analysis), we reduced the number of blocks per window accordingly (4286 consecutive blocks rather than 30,000) to achieve windows of a comparable span.

### Statistical analysis

Permutations were used to test whether differences in *m_e_* between chromosomes were statistically significant. First, a label-switching operation was performed to randomise whether a *B. ino* chromosome was defined as rearranged or non-rearranged, with the rearranged group always consisting of six chromosomes. For each permutation, the differences in mean *m_e_* and barrier window frequency between the randomly defined groups were calculated and used to build null distributions. The observed differences in mean *m_e_* and frequency of barrier windows between rearranged and non-rearranged chromosomes were then compared to these distributions to calculate p-values.

A second permutation test was used to approximate a null distribution for the difference in mean *m_e_* between windows within 1 Mb of rearrangement points, and windows that are elsewhere on rearranged chromosomes. For each permutation, nine points were randomly chosen from rearranged chromosomes and adjacent windows around these points were sampled. The number of adjacent windows sampled for each point was matched to a number of adjacent windows within 1 Mb of a rearrangement point in the real data. Permutations where any window was sampled multiple times were discarded. To avoid under-sampling windows near the ends of chromosomes, adjacent windows were allowed to roll over on to the next rearranged chromosome. The difference in mean *m_e_* between windows adjacent to randomly sampled points and all other windows on rearranged chromosomes, was calculated for each permutation. A total of 100,000 permutations were done to approximate the null distribution. The difference in mean *m_e_* between windows within 1 Mb of rearrangement points and windows that are elsewhere on rearranged chromosomes, was compared to the null distribution to calculate a p-value.

Spearman’s *ρ* was calculated for chromosome length and mean *m_e_*. All analysis were performed in R (R Core Team 2021).

## Data availability

Raw sequencing reads and the *Brenthis daphne* genome assembly are available at the European Nucleotide Archive under project accession PRJEB56310. The scripts VCF_stats.py and min-imap2synteny.py, as well as the R code for carrying out at permutation tests are available at https://github.com/A-J-F-Mackintosh/Mackintosh_et_al_2022_Binodaphne.

## Acknowledgments

We would like to thank Marian Thompson and Robert Foster (both Edinburgh Genomics) for preparing Pacbio and HiC sequencing libraries and Katy MacDonald for help in the molecular lab. We also thank Maria Jesus Cañal Villanueva and Luis Valledor (Universidad de Orviedo) for help with fieldwork logistics, as well as Vlad Dincă, Raluca Vodă, and Sabina Vila for contributing samples. We would like to thank Staffan Bensch and Deborah Charlesworth for insightful comments on an earlier version of the manuscript and Sam Ebdon for helping to improve Figure 1.

