## Supplementary material for "Chromosome fissions and fusions act as barriers to gene flow between *Brenthis* fritillary butterflies"

### Supplementary Materials

#### Supplementary Note 1

The ability to accurately infer demographic parameters from the bSFS depends on a number of variables. A single block only contains information about a single genealogy, so many blocks are required to make accurate inference. The power to infer demographic parameters also depends on the amount of recombination between blocks. However, recombination within blocks can introduce bias, because the analytic calculation for the bSFS assumes no recombination within blocks. So recombination involves a trade-off between power and potential for bias in estimates.

With this in mind, we used gIMble simulate to investigate how recombination affects our ability to estimate demographic parameters in windows across the genome. We simulated windows of equivalent size as those in the real data under the best fitting genome-wide demographic model (Figure 2C). Each simulation contained 30,030 blocks of 64 bases, equivalent to a window of length 45.76 kb split up into 715 blocks. The bSFS was calculated by recording the mutation counts for all 42 possible pairwise comparisons ($6 \times 7$ unphased diploid individuals from *B. ino* and *B. daphne* respectively).

We performed 100 replicate simulations for six different per-base recombination rates: $1 \times 10^{-9}$, $5 \times 10^{-9}$, $9 \times 10^{-9}$, $1.3 \times 10^{-8}$, $1.7 \times 10^{-8}$, $2.1 \times 10^{-8}$. We estimated demographic parameters ($N_e$ and $m_e$) for each replicate, while fixing the split time ($T$), analogous to the the window-wise analysis on the real data. However, we used free optimisation rather than a grid, because the former provides finer parameter estimates. Importantly, it is possible to identify simulation replicates which have not converged to their maximum composite lnCL (MCL), because the parameters they were simulated under are known. This task is much more challenging with the *Brenthis* data (because the parameters are unknown) and so requires the use of a grid.

Comparing results under different recombination rates (Figure S1), we find that there is little power to accurately estimate $m_e$, the parameter we are most interested in, when the recombination rate is $1 \times 10^{-9}$. As recombination increases, estimates of $m_e$ become closer to the true value ($1.811 \times 10^{-7}$). However, $m_e$ is often underestimated at higher recombination rates (Figure S1). For example, the mean estimate of $m_e$ when r $= 2.1 \times 10^{-8}$ is $1.195 \times 10^{-7}$.

This power analysis on simulated data shows that given plausible recombination rates, our

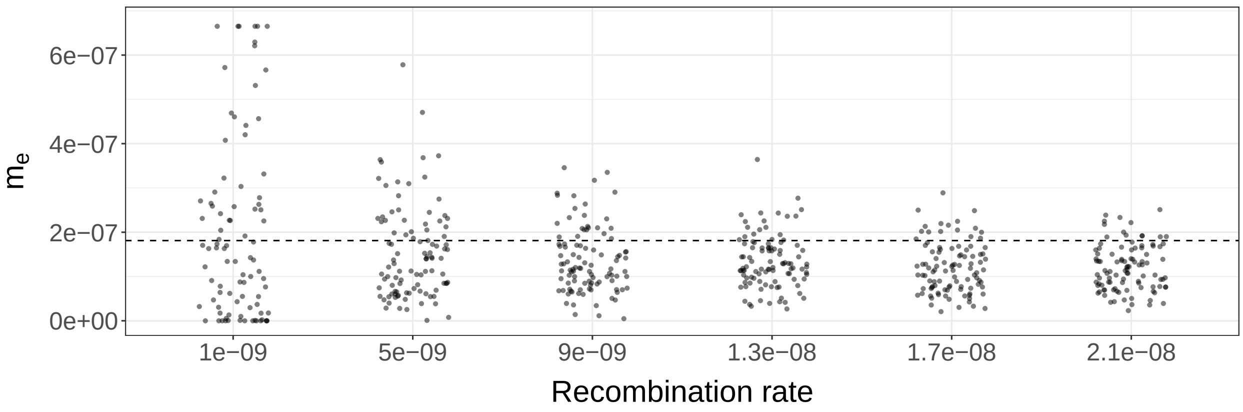


Figure S1: Estimates of effective migration rate ($m_e$) from simulations with different recombination rates. Simulation replicates are plotted as jittered points around the recombination rate that they were simulated under. The simulated $m_e$ ($1.811 \times 10^{-7}$) is plotted as a dashed horizontal line.

analyses of $m_e$ in windows (of 30,000 consecutive blocks) have reasonable power, even though estimates suffer from some downward bias. Although we lack estimates of recombination rate in *B. daphne* and *B. ino*, we expect the mean crossover rate to be approximately $8.5 \times 10^{-9}$ (equivalent to a single crossover per male meiosis for 14 chromosome pairs). In addition, windows in the *Brenthis* dataset often span much greater distances than 45.76 kb (median window span 122 kb) because genic and repetitive regions are removed. This increases the amount of between-block recombination and therefore power.

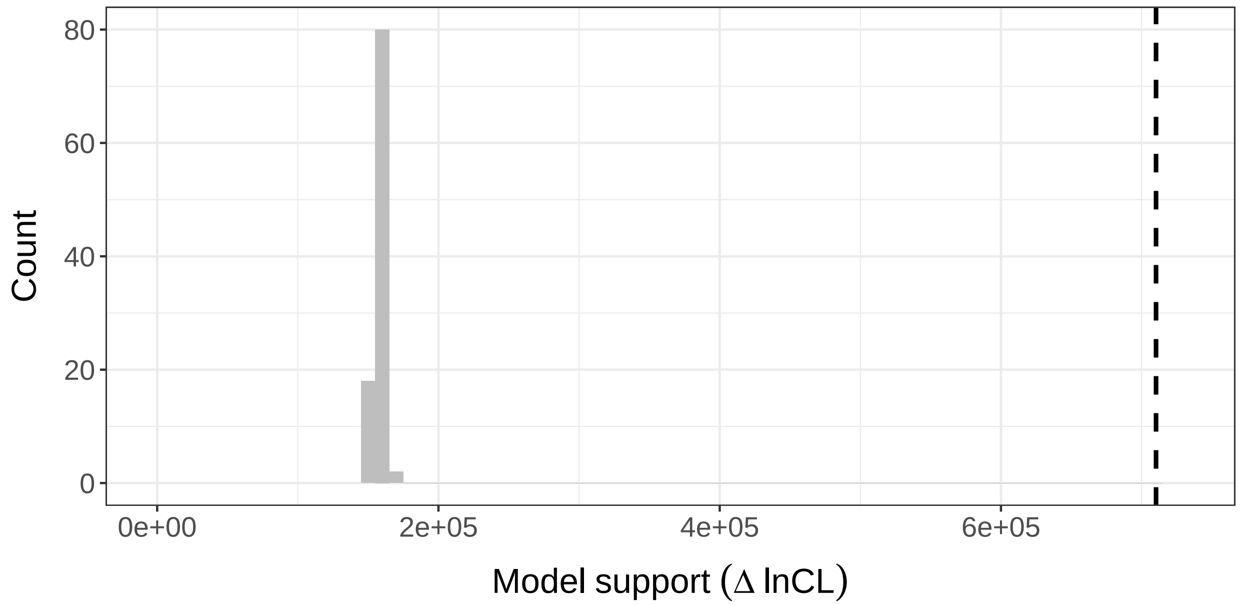


Figure S2: A histogram showing the improvement in model support ($\Delta$ lnCL) between the $DIV$ and $IM_{\rightarrow Bda}$ models for 100 parametric bootstrap replicates, each simulated under the same $DIV$ history. The improvement in fit for the real data is marked with a dashed vertical line..

Table S1: Sampling locations and other metadata for individuals used in this study. In the data column, the source of the data is denoted as TS (This Study) or M2022 (Mackintosh *et al.* 2022).

| **Sample** | **Date** | **Species** | **Sex** | **Locality** | **Region** | **Country** | **Lat** | **Long** | **Collector** | **Data** |
|---|---|---|---|---|---|---|---|---|---|---|
| ES_BD_1141 | 25/7/2018 | daphne | Female | Meranges, La Cerdanya | Catalunya | Spain | 42.435 | 1.797 | RV, Sabina Vila | Pacbio, WGS; TS |
| ES_BD_1489 | 29/7/2009 | daphne | Female | Prioro | Castile and León | Spain | 42.937 | -4.964 | RV | WGS; TS |
| ES_BD_1490 | 24/7/2008 | daphne | Female | Uña | Castile-La Mancha | Spain | 40.231 | -1.960 | RV | WGS; TS |
| FR_BD_1329 | 15/7/2019 | daphne | Female | D620 | Aude | France | 42.997 | 2.053 | KL | WGS, HiC; TS |
| GR_BD_1491 | 26/7/2013 | daphne | Female | Rhodopi, Frakto Forest | Drama | Greece | 41.504 | 24.400 | RV | WGS; TS |
| IT_BD_1493 | 26/6/2012 | daphne | Male | Saguccio, Aspromonte | Aspromonte | Italy | 38.080 | 15.830 | RV | WGS; TS |
| RO_BD_956 | 17/7/2018 | daphne | Male | Pin1000m, Lupsa, Apuseni Mt. | Alba | Romania | 46.416 | 23.192 | KL, AH, DL, RV | WGS; TS |
| ES_BI_364 | 05/07/2017 | ino | Male | Somiedo, Braña de Mumian | Asturias | Spain | 43.068 | -6.24 | KL | WGS, reference genome; M2022 |
| ES_BI_375 | 05/07/2017 | ino | Male | Somiedo, Braña de Mumian | Asturias | Spain | 43.068 | -6.24 | KL | WGS; M2022 |
| FR_BI_1497 | 11/08/2012 | ino | Female | Larche (Les Marmottes) | Alpes-de-Haute-Provence | France | 44.446 | 6.851 | Vlad Dincă, Raluca Vodă | WGS; M2022 |
| RS_BI_1496 | 29/6/2014 | ino | Male | Čeganica | - | Serbia | 43.396 | 22.368 | RV | WGS; TS |
| SE_BI_1495 | 13/7/2016 | ino | Female | Älvsbyn | Norrbotten | Sweden | 65.668 | 20.955 | RV | WGS; TS |
| UA_BI_1494 | 20/6/2014 | ino | Female | Kruglyanka, Novaya Vodolaga | Kharkiv oblast | Ukraine | 49.817 | 35.733 | RV | WGS; TS |
